# MusMorph, a database of standardized mouse morphology data for morphometric meta-analyses

**DOI:** 10.1101/2021.11.11.468142

**Authors:** Jay Devine, Marta Vidal-García, Wei Liu, Amanda Neves, Lucas D. Lo Vercio, Rebecca M. Green, Heather A. Richbourg, Marta Marchini, Colton M. Unger, Audrey C. Nickle, Bethany Radford, Nathan M. Young, Paula N. Gonzalez, Robert E. Schuler, Alejandro Bugacov, Campbell Rolian, Christopher J. Percival, Trevor Williams, Lee Niswander, Anne L. Calof, Arthur D. Lander, Axel Visel, Frank R. Jirik, James M. Cheverud, Ophir Klein, Ramon Y. Birnbaum, Amy E. Merrill, Rebecca R. Ackermann, Daniel Graf, Myriam Hemberger, Wendy Dean, Nils D. Forkert, Stephen A. Murray, Henrik Westerberg, Ralph S. Marcucio, Benedikt Hallgrímsson

**Affiliations:** Alberta Children’s Hospital Research Institute, University of Calgary, Calgary, AB, CANADA; The McCaig Institute for Bone and Joint Health, University of Calgary, Calgary, AB, CANADA; Department of Cell Biology and Anatomy, Cumming School of Medicine, University of Calgary, Calgary, AB, CANADA; Department of Biology, McMaster University, Hamilton, ON, CANADA; School of Dental Medicine, University of Pittsburgh, Pittsburgh, PA, USA; Department of Orthopedics, University of California, San Francisco, San Francisco, CA, USA; Department of Biological Sciences, University of Calgary, Calgary, AB, CANADA; Center for Craniofacial Molecular Biology, Herman Ostrow School of Dentistry, University of Southern California, Los Angeles, CA, USA; Department of Biochemistry and Molecular Medicine, Keck School of Medicine, University of Southern California, Los Angeles, CA, USA; Department of Biochemistry and Molecular Biology, Cumming School of Medicine, University of Calgary, Calgary, AB, CANADA; Institute for Studies in Neuroscience and Complex Systems (ENyS) CONICET, Buenos Aires, ARGENTINA; Information Sciences Institute, Viterbi School of Engineering, University of Southern California, Marina del Rey, CA, USA; Department of Comparative Biology and Experimental Medicine, Faculty of Veterinary Medicine, University of Calgary, Calgary, AB, CANADA; Department of Anthropology, Stony Brook University, New York, NY, USA; Department of Craniofacial Biology, University of Colorado, Aurora, CO, USA; Department of Molecular, Cellular and Developmental Biology, University of Colorado Boulder, Boulder, CO, USA; Department of Anatomy and Neurobiology, University of California, Irvine, Irvine, CA, USA; Center for Complex Biological Systems, University of California, Irvine, Irvine, CA, USA; Environmental Genomics and Systems Biology Division, Lawrence Berkeley National Laboratory, Berkeley, CA, USA; U.S. Department of Energy Joint Genome Institute, Lawrence Berkeley National Laboratory, Berkeley, CA, USA; Department of Biology, Loyola University Chicago, Chicago, IL, USA; Program in Craniofacial Biology and Department of Orofacial Sciences, University of California, San Francisco, San Francisco, CA, USA; Department of Pediatrics and Institute for Human Genetics, University of California, San Francisco, San Francisco, CA, USA; Department of Life Sciences, Faculty of Natural Sciences, The Ben-Gurion University of the Negev, Beer-Sheva, ISRAEL; Department of Archaeology, University of Cape Town, Cape Town, Rondebosch, SOUTH AFRICA; Human Evolution Research Institute, University of Cape Town, Cape Town, Rondebosch, SOUTH AFRICA; School of Dentistry, Faculty of Medicine and Dentistry, University of Alberta, Edmonton, AB, CANADA; Department of Medical Genetics, Faculty of Medicine and Dentistry, University of Alberta, Edmonton, AB, CANADA; Department of Radiology, Cumming School of Medicine, University of Calgary, Calgary, AB, CANADA; The Jackson Laboratory, Bar Harbor, ME, USA; Department of Bioimaging Informatics, MRC Harwell Institute, Oxfordshire, UK; School of Natural Sciences, University of California, Merced, Merced, CA, USA; Department of Developmental and Cell Biology, University of California, Irvine, CA, USA

**Author notes:** Corresponding author: Benedikt Hallgrímsson.

**Keywords:** Mouse, phenomics, craniofacial, imaging pipelines, deep learning, morphometrics, micro-computed tomography, FaceBase

## Abstract

Complex morphological traits are the product of many genes with transient or lasting developmental effects that interact in anatomical context. Mouse models are a key resource for disentangling such effects, because they offer myriad tools for manipulating the genome in a controlled environment. Unfortunately, phenotypic data are often obtained using laboratory-specific protocols, resulting in self-contained datasets that are difficult to relate to one another for larger scale analyses. To enable meta-analyses of morphological variation, particularly in the craniofacial complex and brain, we created MusMorph, a database of standardized mouse morphology data spanning numerous genotypes and developmental stages, including E10.5, E11.5, E14.5, E15.5, E18.5, and adulthood. To standardize data collection, we implemented an atlas-based phenotyping pipeline that combines techniques from image registration, deep learning, and morphometrics. Alongside stage-specific atlases, we provide aligned micro-computed tomography images, dense anatomical landmarks, and segmentations (if available) for each specimen (*N*=10,056). Our workflow is open-source to encourage transparency and reproducible data collection. The MusMorph data and scripts are available on FaceBase (www.facebase.org, doi.org/10.25550/3-HXMC) and GitHub (https://github.com/jaydevine/MusMorph).

## Background & Summary

Understanding how genes, development, and the environment produce variation in complex morphological traits is a core challenge in biology with evolutionary and clinical implications. Explanations for the generation of variation tend to cohere around the genotype-phenotype map concept. Genetic variation and genetic effects, like epistasis and pleiotropy, drive variation in developmental processes that act at different times and scales in anatomical context^1–3^. Specific developmental and genetic mechanisms then operate alongside embedded mechanisms, such as nonlinearities^4,5^ and gene redundancy^6^, to modulate these effects to express a phenotype^7–9^. Despite recent insights into these phenomena, the developmental-genetic basis for morphological variation remains largely unknown, as there are likely many overlapping and coordinated mechanisms involved, each with relative contributions^10^. To help disentangle these mechanisms, it is important to build and integrate large phenotypic databases for model organisms^11–14^. In this work, we present MusMorph, a database of standardized mouse morphology data for meta-analyses of morphological variability and variation, particularly in the craniofacial complex and brain.

The laboratory mouse is a useful model organism for studying the mechanisms of morphological variation because of its 99% genetic homology with humans, short gestation, and rich set of tools for manipulating the genome in a controlled environment. Unfortunately, phenotypic data are often biased by laboratory-specific data collection protocols. The International Mouse Phenotyping Consortium (IMPC, www.mousephenotype.org) was born out of a need to determine the relationship between genotype and phenotype with standardized phenotypic data. Using micro-computed tomography (μCT) and optical projection tomography, the consortium has studied the anatomy of mouse lines heterozygous or homozygous for a single gene mutation, particularly at embryonic day E9.5, E14.5-15.5, and E18.5^15–20^. Less emphasis has been placed on μCT imaging and analysis of adults and mid-gestation (E10 to E11) mutants, where critical developmental events, like fusion of the craniofacial prominences, occur. Mouse lines with normal (non-pathological) levels of variation, such as recombinant inbred strains and outbred strains with high heterozygosity^21–23^, have also been poorly characterized. Quantifying such variation is important, because it drives disease susceptibility and course of disease in humans.

Recently, model organism phenotyping has transitioned from manual linear measurements to fully automated computational pipelines. One common approach is voxel-based morphometry^24,25^. Voxel-based morphometry is based on the analysis of deformation fields obtained via image registration. After spatially aligning images to an average atlas, the deformation fields can be quantitatively compared between groups on a voxel-wise basis to identify differences in morphology. Voxel-based morphometry remains a pillar of shape analysis, because it can localize small regions of shape change without any *a priori* knowledge of the anatomy, but it is prone to the multiple testing problem^26,27^. Another approach is atlas-based geometric morphometrics, which instead uses registration fields to automatically derive landmarks, or Cartesian coordinate points that are homologous across samples. Geometric morphometrics is central to evolutionary biology and developmental biology, among other fields, because landmarks allow for statistically tractable quantifications of morphological variation, as well as intuitive visualizations^28^. These advantages continue to fuel development of novel geometric morphometric pipelines and extensions^29–33^. Yet large-scale morphometric analyses remain rare due to the sparsity of standardized landmark data.

Here, we introduce MusMorph, a database of standardized mouse morphology data generated with an open-source, atlas-based phenotyping pipeline that integrates techniques from image registration, deep learning, and morphometrics. We compiled the database (*N*=10,056) using μCT scans of mice from a variety of strain/genotype combinations and developmental stages, including E10.5, E11.5, E14.5, E15.5, E18.5, and adulthood. Most of MusMorph is composed of head morphology data, but there are also whole-body embryo data for different integrative analyses. We provide (1) a developmental atlas for each timepoint; (2) a rigidly aligned and preprocessed μCT scan, dense anatomical landmarks, and segmentations (if available) for each specimen; (3) a set of scripts for transforming and comparing an input scan to an atlas; (4) an approach to validate the transformed landmark data and optimize it, if needed. To ensure reproducibility and data sharing, we make the data freely accessible from FaceBase^34^ (www.facebase.org, doi.org/10.25550/3-HXMC)^35^ and our code from GitHub (https://github.com/jaydevine/MusMorph). By incorporating substantial developmental and genetic variation alongside a rich set of metadata, MusMorph will enable standardized morphometric analyses of genotype-phenotypes to better understand the mechanistic basis for morphological variation.

## Methods

### Mice

We compiled mouse embryos and adults from numerous sources. The mouse lines for the E15.5 and E18.5 datasets were generated by the IMPC. These mice were produced and maintained on a C57BL/6N genetic background, with support from C57BL/6NJ, C57BL/6NTac or C57BL/6NCrl. More details about husbandry practices can be found at https://www.mousephenotype.org/impress. The mouse lines for the E10.5, E11.5, E14.5, and adult datasets were produced on a variety of genetic backgrounds at different institutions for studies of craniofacial variation. We hereafter refer to these lines as the Calgary mice, because they were ultimately imaged at the University of Calgary. Specific information about study protocols, such as husbandry practices and genotyping, should be gleaned from the MusMorph dataset summaries on FaceBase or the original studies themselves. Each dataset within the MusMorph project on FaceBase represents a study or set of studies defined by a common study design that yielded similar mouse lines. Details about the experimental design were obtained from the original studies listed in the “Publication(s)” section of each dataset. In addition, we provide a supplementary comma-separated values (CSV) file (Study_Metadata.csv) in the project-wide metadata dataset^36^ on FaceBase that lists the associated studies.

### Micro-computed tomography

#### Sample preparation

Each IMPC embryo underwent a hydrogel stabilization protocol^37^ to prepare for diffusible iodine-based contrast-enhanced μCT (diceCT)^38^. This involved incubating the embryo in a hydrogel solution composed of 4% (wt) paraformaldehyde, 4% (wt/vol) acrylamide (Bio-Rad, USA), 0.05% (wt/vol) bis-acrylamide, 0.25% VA044 Initiator (Wako Chemicals, USA), 0.05% (wt/vol) saponin (Sigma-Aldrich, Germany), and phosphate-buffered saline at 4°C for 3 days. Following incubation, the air in the specimen tube was replaced with nitrogen gas and the tube was immersed in a 37°C water bath for 3 h. The whole embryo was then stained with a 0.025 N to 0.1 N Lugol’s iodine (I_2_KI) solution (Sigma-Aldrich, Germany) for 24 h and mounted in agarose for diceCT. This approach has become a popular alternative to magnetic resonance imaging because it is faster, cheaper, and still offers remarkable contrast, allowing for high-throughput phenotyping of soft and hard tissue^38^.

The Calgary embryos were subjected to different fixation and staining protocols. Each embryo acquired prior to 2017 was fixed in a solution of 4% (wt) paraformaldehyde and 5% (wt) glutaraldehyde. The specimen was next submerged in the CystoCon Ray II (iothalamate meglumine) contrast agent for one hour to stain external morphology. Embryos obtained after 2017 were put through a nucleic acid stabilization protocol that allows for examination of RNA in embryos scanned via μCT^39^. Each embryo was fixed with the PAXgene Tissue FIX solution (Qiagen, PreAnalytics, cat #765312), incubated overnight (17 h +/− 1 h) at room temperature, then transferred to a solution of PAXgene Tissue STABILIZER prepared to manufacturer specification (Qiagen, PreAnalytics, cat #765512). For diceCT, each specimen was placed in a solution of PAXgene Tissue STABILIZER and 1% to 3.75% (wt/vol) Lugol’s iodine for 24 h. The head of every embryo was dissected before being mounted in either agarose or soft wax, which was covered by a microcentrifuge tube and infused with 50-100 μl of tissue stabilizer.

Each Calgary adult was set up with a standardized storage and mounting protocol. The mouse carcass was stored at −20°C after euthanasia. Prior to the day of scanning, the mouse was retrieved and thawed overnight at 4°C. The carcasses were then wrapped in foam and placed into a 37 mm diameter sample holder for μCT.

#### Imaging

The IMPC embryos were imaged at six centers, including the Baylor College of Medicine, Czech Center for Phenogenomics, MRC Harwell, Toronto Centre for Phenogenomics, The Jackson Laboratory, and University of California, Davis. A 3-D image of each iodine-stained whole embryo was acquired with a Skyscan 1172 μCT scanner (Bruker, Kontich, Belgium) at 100 kVp and 100 μA. The raw images were initially obtained with isotropic voxels but variable spatial dimensions and resolutions, ranging between 0.002 mm to 0.04 mm. Image projections were reconstructed into a digital stack using the Feldkamp algorithm^40^.

The Calgary mice were imaged in the 3-D Morphometrics Center at the University of Calgary. A 3-D image of each stained embryo head was obtained with either (a) a Scanco μCT 35 scanner (Scanco Medical, Brütisellen, Switzerland) at 45 kV and 177 μA or (b) a ZEISS Xradia Versa 520 X-ray microscope (Carl Zeiss AG, Oberkochen, Germany) at 40-50 kV, 4-5 W, and 2 s exposure time. A 3-D image of each adult skull was acquired with either (a) a Scanco vivaCT 40 μCT scanner (Scanco Medical, Brütisellen, Switzerland), (b) a Scanco vivaCT 80 μCT scanner (Scanco Medical, Brütisellen, Switzerland), or (c) a Skyscan 1173 v1.6 μCT scanner (Bruker, Kontich, Belgium) at 55-80 kV and 60-145 μA. Like the IMPC data, these original images were obtained with isotropic voxels but variable spatial dimensions and resolutions. Embryo image resolutions ranged between 0.007 mm and 0.027 mm, whereas adult resolutions ranged between 0.035 mm and 0.044 mm. Image projections were reconstructed with the integrated Scanco software, the ZEISS XMReconstructor software, or the Skyscan NRecon v1.7.4.2 software.

### Image preprocessing

We preprocessed each image to account for differences in image acquisition that would interfere with the atlas-based registration workflow described below (Fig. 1). The preprocessing scripts are provided in the MusMorph GitHub repository (https://github.com/jaydevine/MusMorph/tree/main/Preprocessing). In this preprocessing step, we first converted the reconstructed imaging data (.nrrd, .aim, .tiff) to the Montreal Neurological Institute (MNI) .mnc format using file conversion scripts written in Bash and Python (see AIM_to_MNC.sh, NII_to_MNC.sh, TIFF_to_MNC.sh, DCM_to_MNC.sh, and NRRD_to_MNC.py). As part of the open-source MINC library (http://bic-mni.github.io/man-pages/), the .mnc format is implemented using HDF5 (Hierarchical Data Format, version 5), which supports hierarchical data structure, internal compression, 64-bit file sizes, and other modern features^41^.

**Figure 1.**
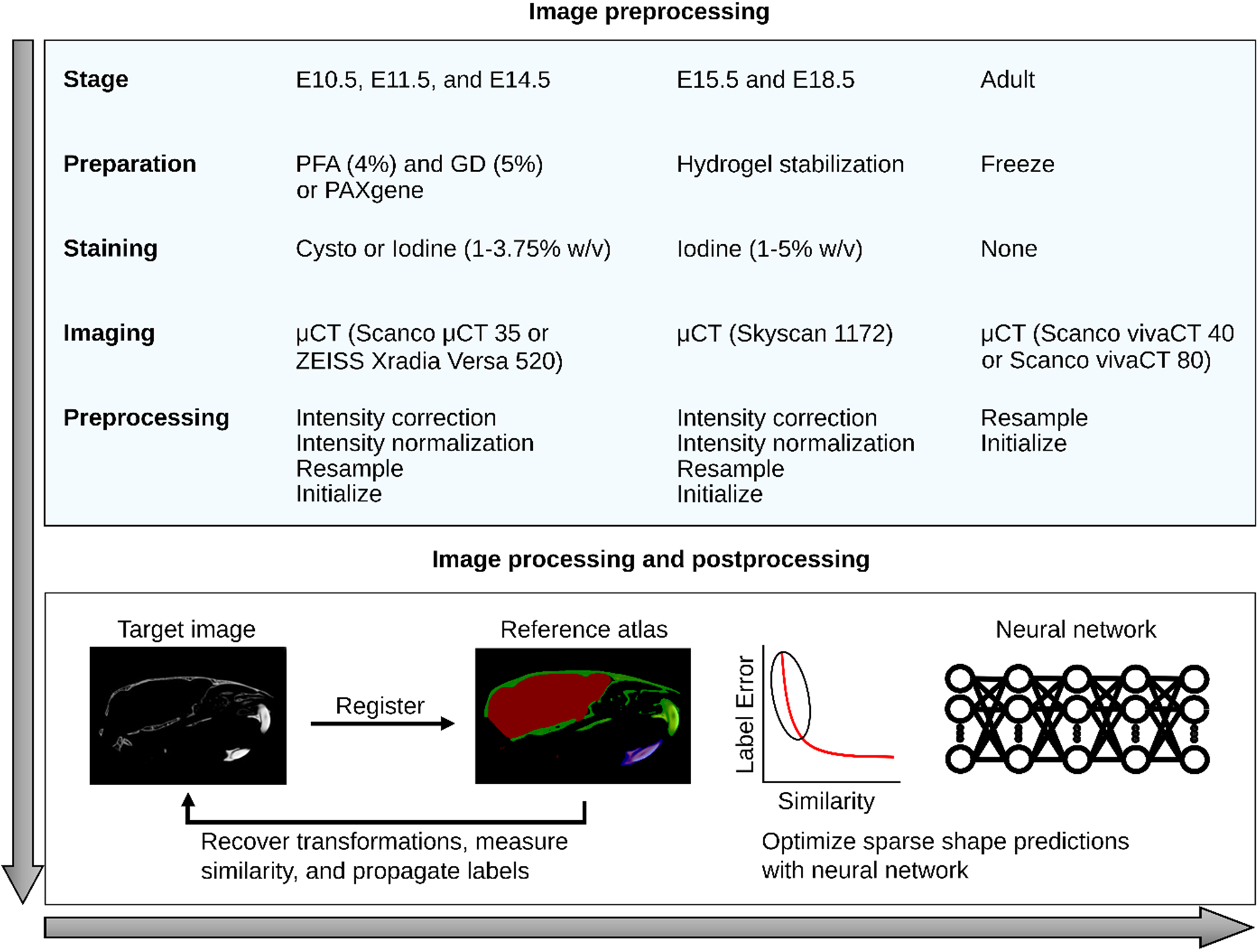
Schematic overview of the phenotyping pipeline. Specimens were staged, prepared (fixed/stored), stained, and imaged with different but standardized lab-specific protocols. While the E10.5, E11.5, E14.5, and adult specimens were obtained in Calgary, the E15.5 and E18.5 specimens were acquired from the IMPC. To account for differences in image acquisition (e.g., intensity artifacts, image resolution and dimensions, and position), each image was subjected to a series of preprocessing steps. Next, each preprocessed image was non-linearly registered to a stage-specific reference atlas with a detailed set of landmarks and segmentations. We recovered deformation fields, landmarks, and segmentations (if available) for each specimen. To optimize the landmark predictions of poorly registered specimens, as measured by cross-correlation similarity, a downstream neural network was used.

Staining artifacts, such as extreme intensity gradients and variable penetrance, can bias the image registration process. To minimize intensity inhomogeneities, we applied the N3 method^42^. Since many of the E15.5 images had background noise, where the stained scanning medium was indistinguishable from the anatomy, we employed a thresholding script in Bash (see Threshold.sh). This script computes a lower anatomical density threshold, masks the voxels above this bound and those in proximity via dilation, and equates all voxels outside the mask to 0. To ensure the image resolutions and dimensions were consistent with the atlas, we implemented an image resampling script in Bash (see Downsample_and_Correct.sh). We also used this script to control for differences in bit depth among scanners by including a min-max normalization, which scaled the embryo intensities between 0 and 1. Table 1 outlines the source of the image data, developmental stage, voxel dimensions, image resolutions, stage-specific sample sizes, and the presence or absence of atlas anatomical labels. Note that the E14.5 images were solely used to create another stage-specific atlas, as they are from a smaller, unpublished dataset.

**Table 1.** Summary of imaging data. Source is where the image was acquired. Stage is the age of the specimen at sacrifice. Anatomy is the labelled and scanned anatomy. X, Y, and Z are the voxel lengths of each atlas axis. Resolution is the isotropic resolution of each scan. N is the sample size, with the number of scans awaiting publication of primary research in parentheses. Landmarks and segmentations indicate the presence (✓) or absence (×) of labels on the stage-specific atlas.

| Source | Stage | Anatomy | X | Y | Z | Resolution (mm) | N | Landmarks | Segmentations |
| --- | --- | --- | --- | --- | --- | --- | --- | --- | --- |
| Calgary | E10.5 | Head | 220 | 295 | 350 | 0.012 | 434 | ✓ | × |
| Calgary | E11.5 | Head | 502 | 503 | 390 | 0.012 | 531 | ✓ | × |
| Calgary | E14.5 | Head;<br>Body | 486 | 567 | 723 | 0.027 | 84<br>(84) | ✓ | × |
| IMPC | E15.5 | Head;<br>Body | 486 | 567 | 723 | 0.027 | 1426 | ✓ | ✓ |
| IMPC | E18.5 | Head;<br>Body | 293 | 414 | 667 | 0.054 | 1657 | ✓ | × |
| Calgary | Adult | Skull | 642 | 586 | 979 | 0.035 | 6000<br>(154) | ✓ | ✓ |

Another essential step to all image registration workflows is the initialization, or a rigid alignment between an image pair. Using initialization scripts written in Bash (see Preprocessing.md) and R (Tag_Combine.R), we rigidly transformed each image to a stage-specific atlas or, if an atlas did not exist, an arbitrary but stage-specific reference image. To determine the rigid transformation matrices, we utilized a manual and automated approach, or a strictly automated approach, depending on anatomical orientation. If the mouse was scanned in a random orientation, we rendered a minimum threshold surface in MINC, then manually placed five homologous three-dimensional (3-D) landmarks at anatomical extrema (e.g., ears, nose, top of the head, and back of the head), resulting in an MNI tag point file (.tag) with landmark coordinates. Next, we concatenated the reference and arbitrary landmark matrices, and minimized their 3-D Euclidean distances via least squares. If the specimen was already roughly aligned to the reference image, we performed an automated, intensity-based rigid alignment using the full registration process outlined below (see the “Image Registration and Label Propagation” section). This intensity-based rigid alignment was also repeated for the manually aligned volumes to ensure consistency. With the rigid transformation matrices, we resampled each image into their stage-specific reference coordinate space using tri-linear interpolation.

### Reference atlases

We generated a population average atlas for each stage, excluding E15.5 and adulthood, by spatially normalizing 25 μCT images of wildtype mice with a group-wise registration workflow^43,44^ (Fig. 2 and 3). A nearly identical workflow was used to create the existing E15.5 and adult atlases. The atlas construction script is available in the MusMorph GitHub (https://github.com/jaydevine/MusMorph/tree/main/Processing) and is written in Python (see HiRes_Atlas.py or LoRes_Atlas.py). This script produces Bash scripts that can be executed automatically and in parallel on a compute cluster to maximize computational efficiency. Without massively parallel computing, the volumetric registrations would need to be performed sequentially, each requiring hours of computation and a large amount of memory. Before executing the workflow, the user must upload the initialized images and registration scripts to a compute cluster. In addition, the user needs to install a MINC Toolkit module onto the cluster via Docker (https://bic-mni.github.io/) or GitHub (https://github.com/BIC-MNI/minc-toolkit-v2), or define a pre-existing module, because the scripts utilize the open-source MINC software. An atlas can also be generated locally, but it will be significantly slower without massively parallel computing.

**Figure 2.**
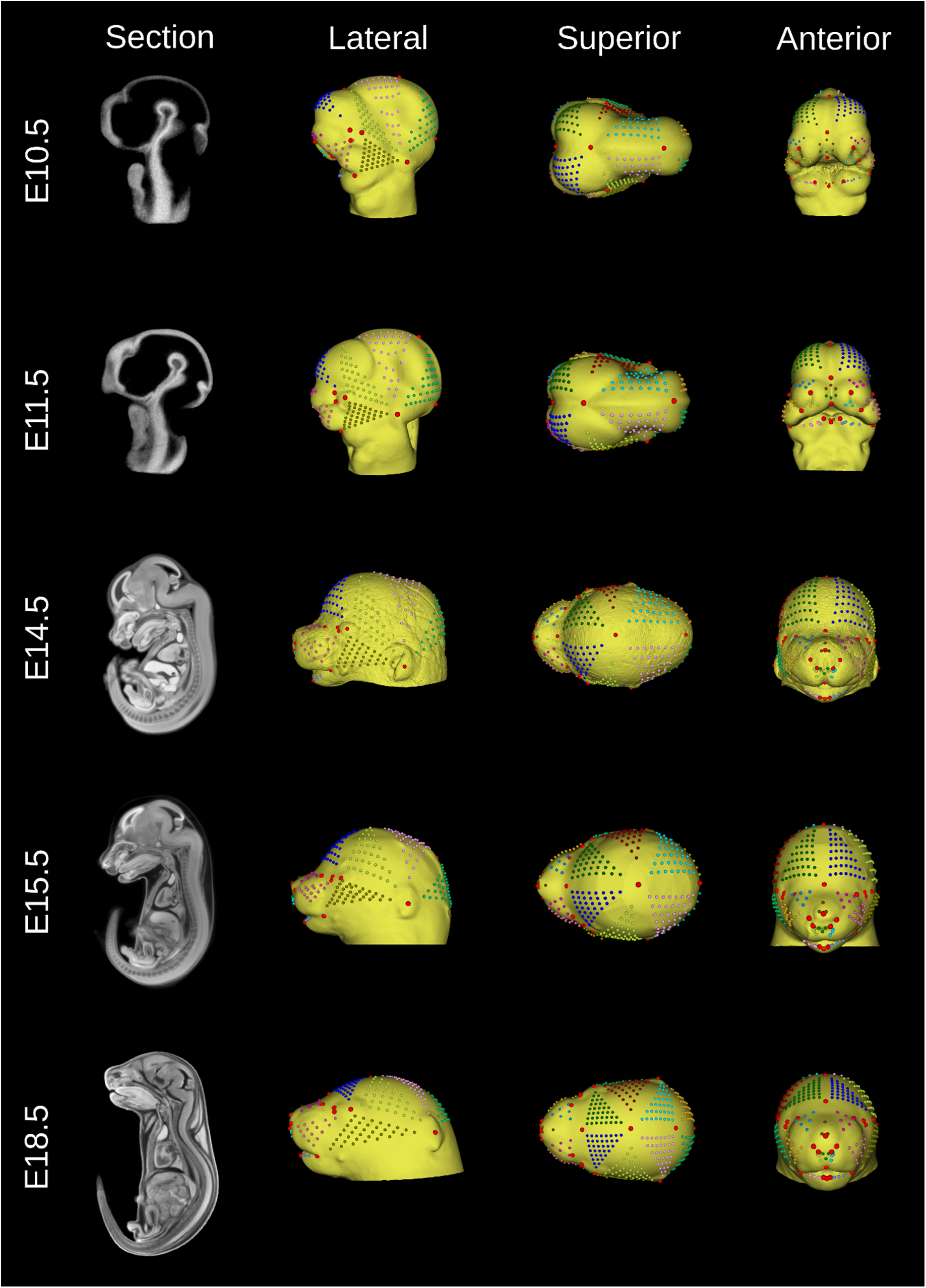
Embryo reference atlases. Sagittal cross-sections of the E10.5 (www.facebase.org/id/6-F00W), E11.5 (www.facebase.org/id/6-F012), E14.5 (www.facebase.org/id/6-F016), E15.5 (www.facebase.org/id/6-F6SE), and E18.5 (www.facebase.org/id/6-F6T4) atlas volumes are shown to display the stained internal anatomy. Each head surface was labelled with a dense landmark configuration to capture global and local aspects of morphology. Lateral, superior, and anterior views of each head isosurface are shown. The semi-landmark patches (small, color-coded points) were interpolated between a set of sparse homologous landmarks (large, red points). They can be slid and resampled for morphometric analyses.

**Figure 3.**
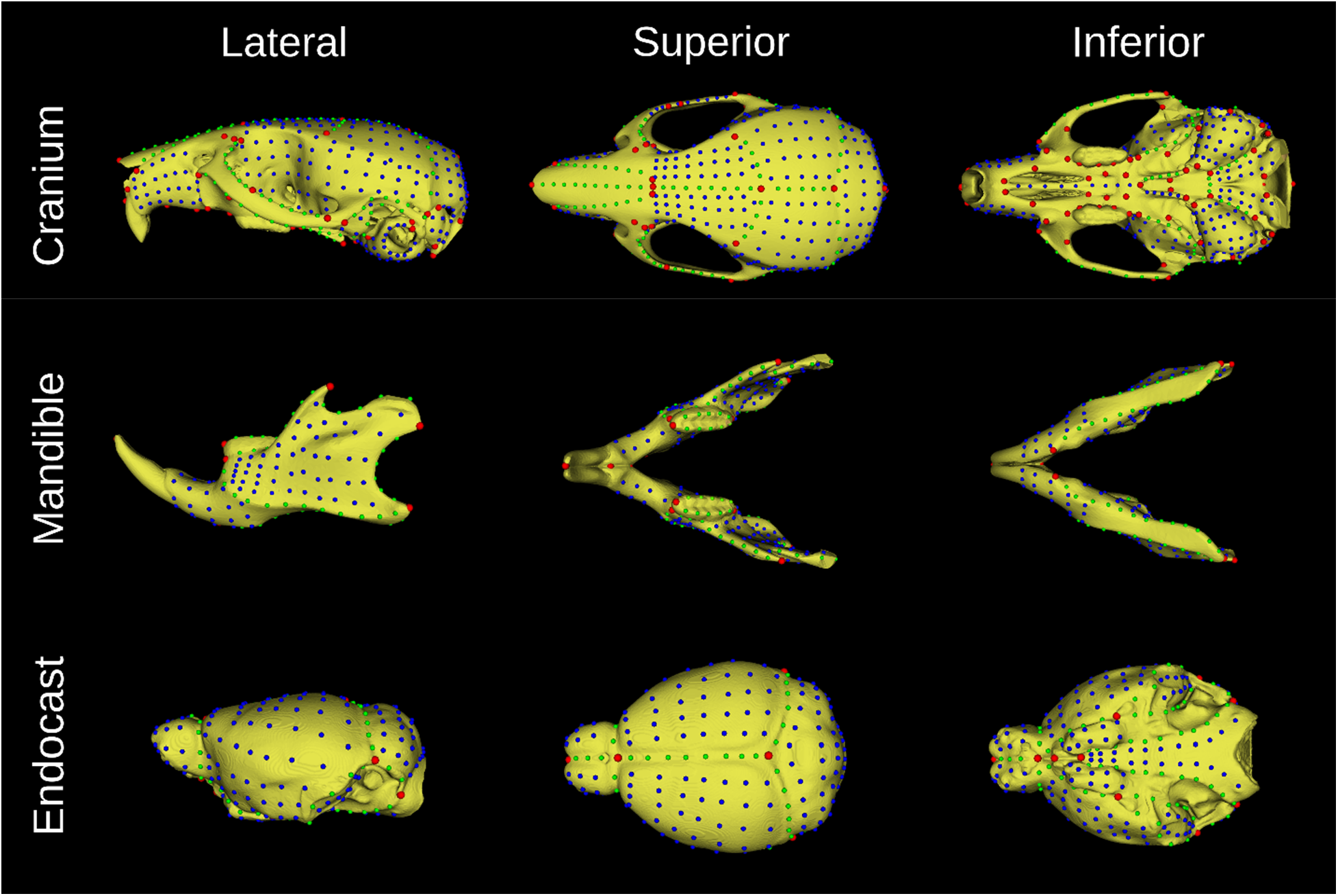
Adult reference atlas. Cranium (top), mandible (middle), and endocast (bottom) surfaces were segmented from the skull atlas (www.facebase.org/id/6-F6VC), then labelled with a dense landmark configuration to capture global and local aspects of morphology. Lateral, superior, and anterior views of each segmentation isosurface are shown. There are sparse landmarks (red) as well as surface (blue) and curve (green) semi-landmarks that can be slid and resampled for morphometric analyses.

Spatial normalization involves an initial affine transformation for global alignment, followed by a deformable transformation for non-linear alignment. To account for global variation in location, orientation, and scale, we computed a series of multi-resolution (coarse to fine) affine transformations among the images by optimizing a cross-correlation objective function^45^. Given that sample-wide pairwise registrations yield an improved affine template^46^, or intensity average, we completed all possible (*N*=25*24) pairwise affine registrations, then averaged the resulting transformation for each specimen. Using the averaged transformations, we resampled each initialized image into the affine coordinate space with tri-linear interpolation and averaged the resulting images to produce an affine template. To correct for local variation in shape, we computed a series of multi-resolution non-linear transformations with the ANIMAL (Automatic Nonlinear Image Matching and Anatomical Labelling) algorithm^47^, again optimizing for cross-correlation. This iterative, four-step process involves non-linearly deforming each mouse to an evolving template at increasingly higher resolutions, with the first template being the affine average and the next three being improved versions of the non-linear average^48^. The final product is a stage-specific average with excellent contrast and a high signal-to-noise ratio.

Since the goal of MusMorph was to aggregate landmark data for morphometrics, and our primary imaging data are head scans, we focused on labelling each atlas head surface with a standardized landmark configuration (Fig. 2 and 3). Specific information about the number of landmarks and their anatomical definitions can be found below in the “Data Records: Landmarks” section. To generate the landmarks, we first rendered a minimum density isosurface in MINC, which uses ITK’s marching cubes algorithm, and saved the 3-D rendering as a Stanford PLY (.ply) file. We then used 3D Slicer^49^ or the MINC Toolkit to acquire a landmark configuration on each surface that provided a comprehensive representation of shape^50^. For the embryos, we used 3D Slicer and the SlicerMorph extension^32^ to identify sparse landmarks and interpolate landmark patches of variable density in between, depending on the size of the area, resulting in dense coverage of the head. This also ensured that the patches were homologous, allowing for a developmental morphospace into which all specimens may be superimposed. Because developmental homology was not a consideration for the adults, we landmarked the adult atlas in MINC using built-in display tools, again ensuring sparse and dense landmark coverage.

Shared developmental pathways lead to correlated morphological variation, or morphological integration^51–57^. To enable analyses of integration, we added landmark configurations to segmented surfaces of the adult skull atlas. We manually segmented the cranium, mandible, and neurocranial endocast (i.e., a proxy for the brain) in MINC, then rendered these segmentations as isosurfaces before landmarking them with a dense configuration. Once again, the landmark details are described below in the “Data Records: Landmarks” section. The segmentations may further be used for surface-based analyses^58^, measures of size (e.g., volume or surface), or as masks to reduce the shape dimensionality of a voxel-based morphometry analysis. Unlike the adult atlas, the embryo atlases do not come with segmentations due to the scope of this work, apart from the pre-existing E15.5 atlas, which has 48 manually segmented structures (http://www.mouseimaging.ca/technologies/mouse_atlas/mouse_embryo_atlas.html).

### Image registration and label propagation

We pairwise registered each image to their stage-specific atlas to obtain a composite (affine and non-linear) transformation for label propagation (Fig. 1). Like the atlas workflow described above, the registration scripts are available in the MusMorph GitHub (https://github.com/jaydevine/MusMorph/tree/main/Processing) and are written in Python (see HiRes_Pairwise.py or LoRes_Pairwise.py). The purpose once more is to produce Bash scripts *en masse* for massively parallel computing on a compute cluster due to the computational requirements of volumetric deformable registration and anatomical labelling. Only the initialized images and registration scripts need to be uploaded to the cluster to execute the workflow. While the pairwise registrations involved the same multi-resolution affine alignment described above, the non-linear alignment differed. Here, we implemented the geodesic SyN (Symmetric Normalization) algorithm^59^, because it was previously validated for atlas-based landmarking and morphometrics of mouse models^44^. The SyN registrations were optimized using cross-correlation. After registration, we used labelling scripts written in Bash and produced via Python (see Label_Propagation.py) to recover the non-linear transformations, concatenate them with the affine transformations, invert them, and propagate the atlas labels to the rigid space of each image.

### Neural network shape optimization

Although top-performing registration algorithms provide an effective and generalizable way to automatically label anatomy, there are instances where outliers and problematic landmarks can alter shape representations. This is particularly true for model organisms, where mutant phenotypes may show little to no resemblance with an atlas. To demonstrate how biological signal can be restored, we implemented a supervised deep learning workflow available in the MusMorph GitHub (https://github.com/jaydevine/MusMorph/tree/main/Postprocessing), which employs scripts written in R and Julia (see GPA_and_Projection.R and Landmark_Optimization.jl)^60^. Using a subset of 68 sparse adult craniofacial landmarks (*N*=2,000) described in previous work^61–65^, we trained a deep feedforward neural network to learn a domain-specific loss function that minimizes automated and manual shape differences. The sparse landmark numbers amenable to optimization (see Optimization_Order.csv)^36^ are available on FaceBase. We focused on the adults because that was the only stage with a large existing set of homologous manual landmarks for training.

We tested the network predictions on a random subset (*N*=500) of adult skulls described further in the “Technical Validation” section. To help others initialize the network without having to retrain it, we provide the adult network model (Calgary_Adult_Cranium_Model.bson) and weights (Calgary_Adult_Cranium_Weights.bson) in the Binary JSON (.bson) file format on GitHub. We also make available the optimized sparse shape predictions for the entire adult crania dataset (Adult_Cranium_Sparse_Landmarks.csv)^36^. Although we focused on adults, this optimization strategy is generalizable, so other research groups with manual landmark data on any structure of the atlases may use the network architecture to improve outlier predictions.

## Data Records

### Specimen metadata

Each specimen is associated with a rich set of identifiers to accommodate morphometric analyses using multiple factors and/or covariates. Alongside detailed metadata descriptions in FaceBase, we provide the specimen metadata as a supplementary CSV file (MusMorph_Metadata.csv)^36^ for convenience and to include auxiliary fields. Table 2 enumerates the metadata and Table S1 summarizes the metadata distributions for each dataset on FaceBase.

**Table 2.** Summary of metadata identifiers.

| Identifier | Description |
| --- | --- |
| Biosample | The name of the specimen, which corresponds to the image and label names. |
| Strain | The background strain of the specimen. |
| Strain_MGI_ID | The MGI ID for the strain. |
| Strain_Type | An attribute of strain that describes whether it is inbred or outbred and lab-derived or wild-derived. |
| Gene | The gene symbol as provided by MGI. |
| Gene_MGI_ID | The MGI ID for the gene. |
| Zygosity | Whether the specimen is homozygous, heterozygous, wildtype, or otherwise (e.g., flox/null) for a given gene mutation. |
| Genotype | A concatenation of the gene symbol and zygosity symbol. |
| Anatomy | The region of anatomy that has been scanned and labelled. |
| Treatment | An environmental effect that the specimen has been treated with. |
| Experimental Group | An identifier derived from genotype that denotes whether the specimen is a control or mutant. |
| Sex | The sex of the specimen. |
| Stage | The age of the specimen in days, either embryonic (E) or postnatal (PN). |
| Life_Phase | An identifier derived from stage that indicates life phase (e.g., gestation vs. adulthood). |
| Dataset | The published or unpublished study (see Study_Metadata.csv) the sample is associated with. |
| Availability | Whether the images and phenotypic data are available or pending publication of a primary research article. |

Fig. 4a-b illustrates the distributions of sex, strain type, and genotype across the embryo and adult datasets. Sex is well-annotated for the E15.5, E18.5, and adult datasets, but is missing (“NA”) for many of the E10.5 and E11.5 specimens. While most of the embryo mouse models were produced on an isogenic inbred background, particularly C57BL/6N, strain diversity is a focal point of the adult datasets. Among the nine adult strain types provided, there are 98 unique background strains. The majority are recombinant inbred lines (e.g., the Collaborative Cross dataset^66^), wild-derived crosses (e.g., the Hybrid dataset^67^), and outbred lines (e.g., the Diversity Outbred dataset^68^). We have included 459 unique genotypes for the embryo datasets, most of which derive from the IMPC dataset^69^, as well as 179 genotypes for the adult datasets. A minority of specimens, including several embryos in the Ap2^70^, B9d^71^, and Bulgy^72^ datasets as well as a few adults in the Brain-Face^73^ dataset, have unknown genotypes (e.g., “−/−;NA” and “+/−;NA” in double knockout designs or “NA” and “+/+ or +/−” in single knockouts) due to genotyping complications in the past. Specimens homozygous for a single gene mutation predominate the embryo datasets, whereas normal wildtype variants comprise the bulk of the adult datasets. Fig. 4c shows the developmental stages represented in MusMorph. Of the 10,056 specimens processed, 40% are embryos and 60% are adults, many of which have just finished maturing around postnatal day 90. All specimens without a recorded stage (“NA”) are mature adults.

**Figure 4.**
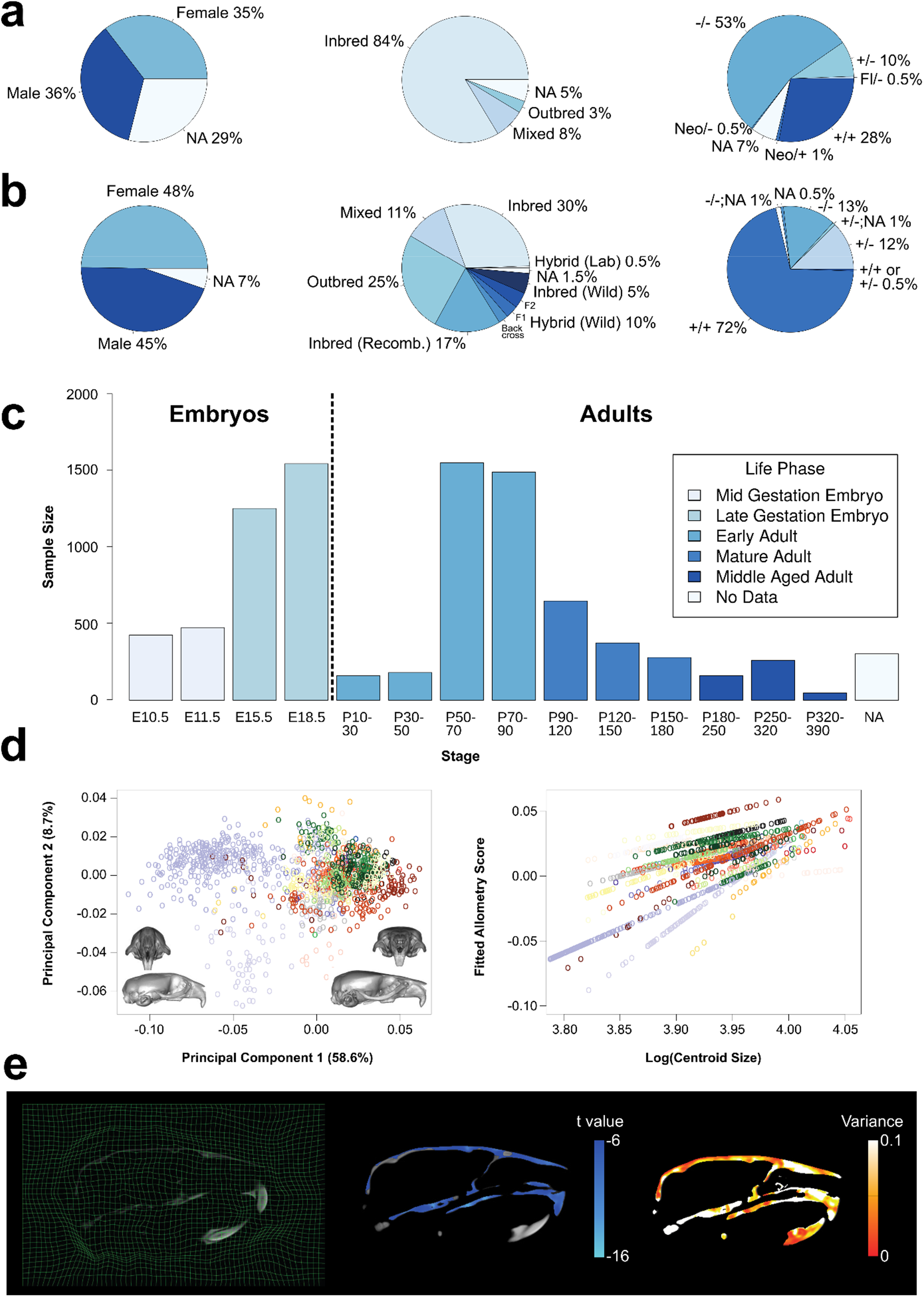
Summary of metadata. (a) Distribution of sex, strain type, and genotype for the embryo datasets. (b) Distribution of sex, strain type, and genotype for the adult dataset. (c) Sample sizes of each developmental stage included in the database. All “NA” specimens are mature or middle-aged adults. (d) Left: Example landmarks and segmentations of the adult skull and endocast (brain). Middle/Right: Morphological analyses, such as PCA and allometry regressions, that one might perform with a dense landmark dataset. Each color in the plot represents a different mouse genotype. (e) Left: Slice visualization of a non-linear deformation grid. Middle/Right: Morphological analyses, such as statistical parametric mapping, that one might perform with a deformation field. The t values show significant (p < 0.05) voxel-wise differences in form (i.e., volume shrinkage) in Ghrhr homozygous mutants relative to wild type, whereas the variance heatmap shows voxel-wise variances in Ghrhr mutants.

It is often desirable to compare mutants to their wildtype counterparts from the same sample because background strains vary. To preserve sample provenance where possible, specimens that are wildtype for a given mutation will have the same gene symbol as their heterozygote and homozygote littermates. For wildtype specimens without litter information, like the IMPC dataset, their genotypes are equated to background strain. Mouse strain nomenclature follows the MGI guidelines, except when the strain design is unknown and has no MGI ID (e.g., novel hybrid backcrosses). We also abbreviate genotypes for complex strain designs using MGI synonyms if available. Furthermore, while most wildtype specimens fall within the control experimental group, there are cases where they can exhibit mutant-like phenotypes and be categorized as such. One example in MusMorph is the artificial selection Longshanks dataset^74^, which through many generations of artificial selection produced wildtype specimens with extreme tibia and craniofacial phenotypes^75,76^.

We selected the above identifiers, because they tend to explain a significant amount of morphological variation in morphometric analyses. For instance, many structures in the mouse are sexually dimorphic, including the shape of the brain^77^ and craniofacial complex^78^, cortical bone size and strength^79^, adipose tissue distribution^80^, and feto-placental growth^81,82^, to name a few. It is also known that classical laboratory strains, such as those in the Strain Comparison dataset^83^, exhibit naturally occurring craniofacial phenotypes^84^. Moreover, gene mutations can interact with a background strain via epistasis to produce different phenotypes^85–87^, like those in the Spry dataset^88^. Another key driver of variation is developmental stage, as differences in age often define a principal axis of allometric variation via correlations with size and/or shape^89–93^. Given the ubiquity of allometry, these correlations can be found across most MusMorph datasets (Fig. 4d). Finally, numerous studies have reported the phenotypic outcomes of single gene mutations, environmental perturbations, and how zygosity modulates these effects^94–96^. These identifiers have corresponding images, landmarks, segmentations, and deformation fields for morphological analyses (Fig. 4d-e).

### Images

We provide the atlases and initialized images for each specimen in the MNI .mnc format. The naming convention for the atlas volumes is *<Source>*_*<Stage>*_*<Anatomy>*_Atlas.mnc. They are categorized as “Imaging Data” in the project-wide dataset^36^ on FaceBase. The naming convention for the initialized volumes is *<Biosample>*.mnc, where *Biosample* is the name of the specimen in the metadata (see the “Specimen metadata” section). One exception is the naming convention for the subset of thresholded E15.5 images, which is *<Biosample>*_Thresh.mnc. These volumes are also categorized as “Imaging Data” across the MusMorph datasets on FaceBase. Each .mnc file has four key attributes: 1) a *named dimension* (*xspace, yspace*, *zspace*), 2) *length* (number of voxels on each dimension), *step* (resolution), and *start* (origin). MINC defines a voxel and world coordinate system, so one can move between them with the simple “voxeltoworld” and “worldtovoxel” MINC commands. If users want to convert between .mnc and different file formats (e.g., raw data, DICOM, NIfTI, Analyze, ECAT, TIFF, Concorde, VFF), there are a variety of other Bash commands available (http://bic-mni.github.io/man-pages/). While the raw IMPC images are freely accessible in the NRRD (.nrrd) format at https://www.mousephenotype.org/data/embryo, the raw Calgary images are available upon request in the AIM (.aim) or TIFF (.tiff) formats.

### Transformations

For each pairwise registration, we recovered an inverted non-linear and composite (affine and non-linear) transformation. Given the file sizes of the non-linear deformation fields (~3 GB on average × 10,000 = 30 TB), we make the transformations available upon request. The deformation fields and composite transformations are in the MNI .mnc and .xfm formats. Each .mnc file shares the same image attributes described above with an additional *named dimension* called *vector_dimension* which describes the non-linear displacement vectors. Each .xfm file contains a header and affine transformation matrix. The naming convention for the deformation fields is *<Biosample>*_ANTS_nl_inverse_grid_0.mnc and *<Biosample>*_ANTS_nl_inverse.xfm, whereas the composite transformations are called *<Biosample>*_origtoANTSnl_grid_0.mnc and *<Biosample>*_origtoANTSnl.xfm. “ANTS” denotes the algorithm and “nl” stands for “non-linear”. Much like the images, the transformations for the subset of thresholded E15.5 volumes have “Thresh” appended to the *<Biosample>* name.

Non-linear deformation fields describe the displacements of each target image voxel to each reference image voxel^97^. By calculating the Jacobian determinant J for every point p (x, y, z) in the deformation field,

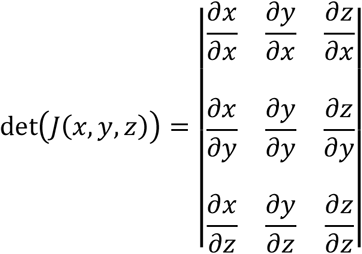

one can quantify the magnitude of morphological change at each voxel (Fig. 4e). A Jacobian determinant of 1 indicates no volume change, whereas determinants greater than 1 indicate volume expansion and determinants between 0 and 1 indicate volume shrinkage. These determinants can also be scaled and sheared with a composite transformation to examine voxel-wise differences in form. Jacobian determinants can be analyzed with voxel-wise tests, such as an ANOVA with a false-discovery rate correction, to map statistics onto the anatomy, a technique otherwise known as statistical parametric mapping (see VBM_Example.R). For example, in Fig. 4e, we use the *RMINC* R package (https://github.com/Mouse-Imaging-Centre/RMINC) to show significant voxel-wise changes (shrinkages) in form between *Ghrhr* mutants^98^ and wildtype specimens, as well as voxel-wise variances in form associated with this mutation.

### Landmarks

We labelled each atlas, and thus every registered mouse embryo and adult, with a standardized landmark configuration (Fig. 2 and 3). The atlas landmark files are named *<Source>*_*<Stage>*_*<Anatomy>*_Atlas_Landmarks.tag. They are stored as “Imaging Data” alongside the atlas volumes on FaceBase^36^. The individual specimen landmark files are named *<Biosample>*_*<Anatomy>*_Landmarks.tag and are similarly categorized as “Imaging Data” across FaceBase. The MNI .tag file format is an ASCII file which stores the coordinates of each landmark in the millimetric world space of the volume. Each .tag file has a header above an array of *p* landmarks (rows) in *k* dimensions (columns). These files can be imported into R individually or collectively as a 3-D array using the *tag2array* function in the custom *morpho.tools.GM* package^99^. Alternatively, the user can employ the *read.csv* function in R to import a vectorized .csv file. We provide landmark .csv files for every developmental stage and anatomical region^36^, each of which contains a matrix of *n* specimens (rows) and *p* × *k* landmark coordinate dimensions (columns). Importantly, there are dense semi-landmarks and sparse fixed landmarks for local and global geometric morphometric analyses of craniofacial, endocast (brain), and mandible morphology. In Fig. 4d, for instance, we show craniofacial shape morphs along the first principal component (PC) in an adult subsample, as well as allometry regressions which relate craniofacial shape to size.

The embryo landmarks are homologous across stages. Table S2 describes the sparse embryo landmarks and their biological definitions. Table S3 lists the embryo semi-landmark patches and their density, both of which are based on the sparse landmarks. The stage-specific semi-landmark patch files can also be found as tab-separated value (TSV) files on GitHub (https://github.com/jaydevine/MusMorph/tree/main/Postprocessing). Each embryo has 22 sparse homologous landmarks within their larger dense configuration. To perform a sparse landmark shape analysis, users may subset the first 22 rows of each 3-D array. Since there are three additional sparse landmarks for the E15.5 and E18.5 specimens, rows 23 to 25 may be included for stage-specific analyses or excluded for ontogenetic analyses.

The adult landmarks are simply homologous within stage. Tables S4, S5, and S6 describe the sparse adult craniofacial, endocast, and mandible landmarks, respectively, as well as their biological definitions. While the adult curve semi-landmarks and surface semi-landmarks are not patch based, they can be slid and resampled using the R scripts on GitHub (see Calgary_Adult_Cranium_Sliding_Semis.R, Calgary_Adult_Mandible_Sliding_Semis.R, and Calgary_Adult_Endocast_Sliding_Semis.R) to mimic patches or any other structure. Much like the embryos, the sparse landmarks are the first 93, 12, and 19 rows of the cranium, endocast, and mandible 3-D arrays, respectively, and can be partitioned for a sparse shape analysis. If users want to generate new landmarks, such as internal landmarks or whole-body landmarks, they can use a script (see Label_Propagation.py), the inverted composite transformations (see the “Transformations” section), and a local or remote compute cluster to propagate the landmarks to an initialized image. To promote standardization, we encourage users to add new landmark subsets to the pre-existing configurations.

### Segmentations

We provide segmentation labels for the E15.5 and adult atlases and specimens to support alternative morphological analyses, such as 3-D visualizations, voxel-based morphometry, volumetric size comparisons, and surface-based image processing pipelines. Other stages do not have segmentation labels due to the scope of this work. The segmentations follow the same naming conventions described above: *<Source>*_*<Stage>*_Atlas_Segs.mnc and *<Biosample>*_Segs.mnc. The atlas segmentations are available as “Imaging Data” on FaceBase^36^, as are the individual segmentation files across various MusMorph datasets. The published E15.5 atlas contains 48 whole body segmentations (http://www.mouseimaging.ca/technologies/mouse_atlas/mouse_embryo_atlas.html)^48^, while the adult atlas comes with cranium, endocast, and mandible segmentations. Each label file is a .mnc volume of integers that matches the dimensionality of the image. To visualize the adult segmentations, for example, the user may load the atlas and label files together and input an integer of 1 to render the endocast, 2 for the cranium, and 3 for the mandible. As with new landmarks, there is the potential to resample new atlas segmentation labels into the initialized space of any image using the composite transformations (see the “Transformations” section) and a local or remote compute cluster (see Label_Propagation.py).

## Technical Validation

### Cross-correlation and root mean squared error

We computed intensity-based, pairwise registrations between each target image (*I*) and a reference atlas (*J*) by optimizing a normalized cross-correlation (NCC) similarity metric:

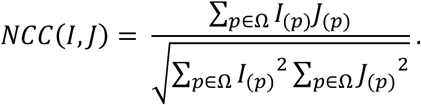

Normalized cross-correlation is calculated for all voxel positions *p* over a discrete domain (*p* ∈ *Ω*). If the domain is the entire 3-D volume and *NCC(I, J)* = 1, the deformed target image and reference image are perfectly aligned. To assess the quality of each registration, we recorded the normalized cross-correlation between each deformed target image and the atlas using code in the labelling scripts (see Label_Propagation.py). Unfortunately, it is difficult to know whether the final registration correlations are “good” or “bad” without relating them to the quality of the labels collected. We investigated the relationship between landmark root mean squared error and cross-correlation in the adult crania training set above to build a quality assessment model. Letting x_ℓ_^(*I*)^ and 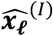 denote the observed (manual) and predicted (automated) Euclidean vectors at landmark ℓ for a target image *I*, the root mean squared error for *p* landmarks is defined as

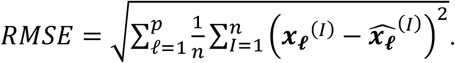

After computing the root mean squared error for each specimen, we regressed these values on their corresponding cross-correlation values with linear, squared, and cubic cross-correlation terms (Fig. 5a). We found a statistically significant non-linear relationship (*R^2^* = 0.3, p < 0.001), such that cross-correlation values below 0.90 resulted in exponentially higher landmark errors. The average root mean squared error was 0.23 mm (95% CI ± 0.002 mm). This mean error is comparable to manual landmark intra-observer detection errors across the skull, which tend to be 0.25 mm or less^44,50^. To verify registration quality across the rest of the database, we calculated cross-correlations for all specimens and stages. The mean cross-correlation values and their standard deviations for E10.5, E11.5, E15.5, E18.5, and adulthood were 0.94 ± 0.07, 0.96 ± 0.04, 0.93 ± 0.02, 0.93 ± 0.12, and 0.95 ± 0.02, respectively (Fig. 5a). These values are on par or higher than those reported in previous mouse registration studies^100^ and speak to the reproducibility of this approach for analyzing variable morphology.

**Figure 5.**
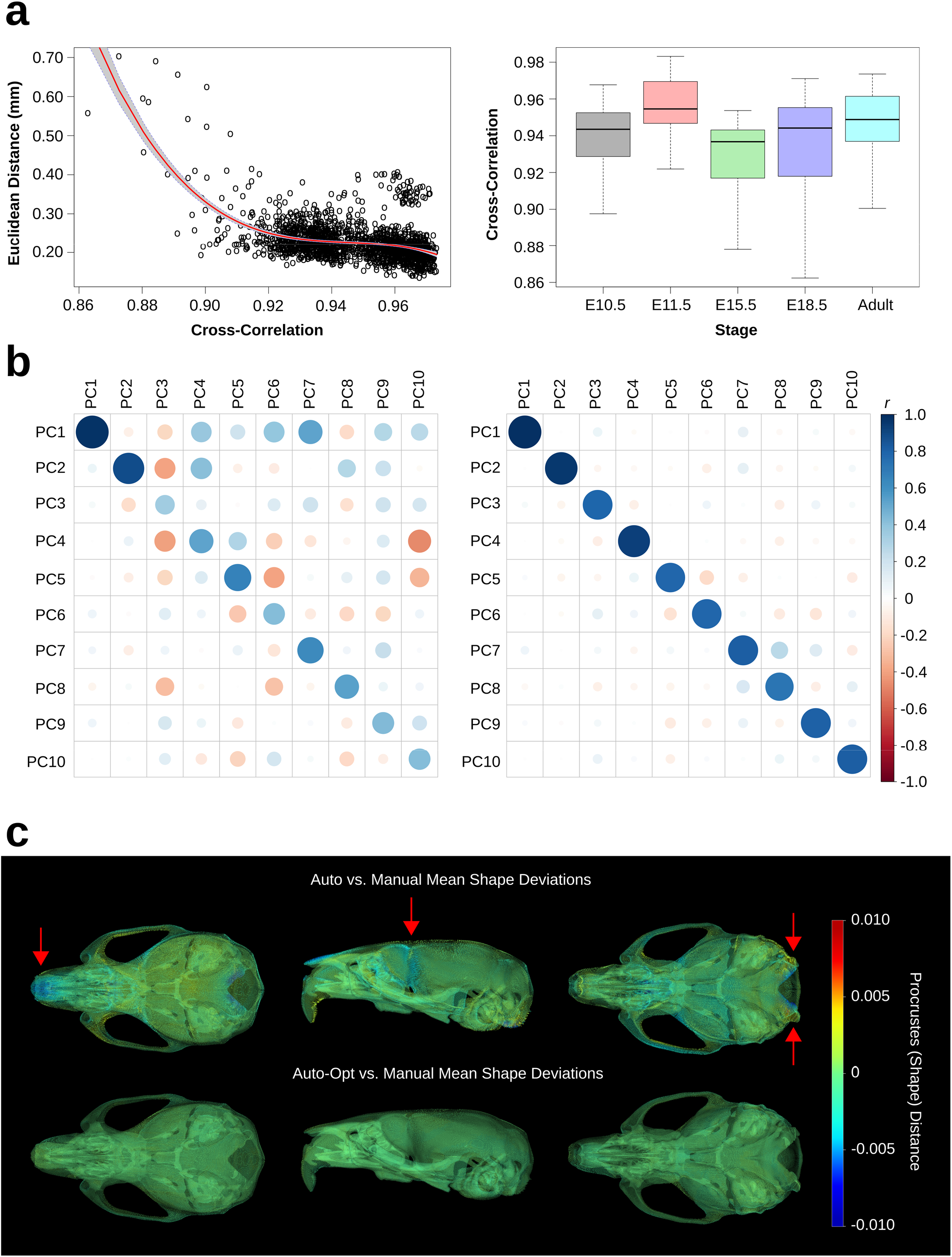
Validation of adult crania test set. (a) Left: Regression of automated-manual Euclidean distances (error) on cross-correlation, a measure of the final target-reference image similarity. Right: Boxplots showing the distribution of cross-correlation values within each developmental stage. (b) Correlation of automated and manual PC scores. Left: Baseline automated PC correlations. Right: Optimized automated PC correlations. (c) Mean shape deviations between the automated and manual datasets. Red arrows indicate error prone areas.

### Covariance patterns and the mean shape

We quantified differences in covariance structure and the sample mean shape between our baseline automated landmarks, the optimized neural network landmarks, and the manual landmarks. To analyze covariance similarity, we projected the automated configurations into the manual PC space and correlated the uncentered PC scores. Fig. 5b shows automated and manual correlations for the first 10 PCs (65.1% of the total variance). The average correlation within PCs for the baseline automated configurations was *r* = 0.6. This measure is biased downwards by lower order automated PCs, which tend to capture residual covariance of the first manual PC. The average correlation within PCs for the optimized automated configurations was *r* = 0.8, suggesting a restoration of signal among the major PCs.

To analyze mean shape deviations, we computed the grand mean shape for the manual landmarks and deformed it to the automated mean shapes via thin-plate spline. We then used the *Morpho* package^101^ in R to generate a deformation heatmap of Procrustes distances at every vertex of the deformed mesh (Fig. 5c). Procrustes distance is equivalent to the root mean squared error between two configurations in shape space. The total distance between the baseline automated mean and manual mean was 0.05, whereas the distance between the optimized automated mean and manual mean was 0.01. Visually, the baseline automated mean shape is largely indistinguishable from the manual mean shape, apart from several known problematic areas^42^. First, the anterior extent of the frontonasal prominence is underestimated. Second, the shape of the foramen magnum is altered. Third, the lateral extent of the frontal bone is underestimated, likely because there are no sparse landmarks to interpolate there; however, this area is well-covered by the dense landmark configurations. Optimization successfully corrected errors at these problematic locations.

### Outliers and stage-specific shape distributions

For each stage, we calculated the Procrustes distance between the mean shape and every configuration to obtain shape distributions and identify outliers (Fig. S1). We defined outlier shapes as those with a Procrustes distance above *Q*_3_ + 1.5 × *IQR*, where *Q*_3_ is the third quartile and *IQR* is the interquartile range. Next, we displayed a minimum threshold isosurface of each outlier image alongside its landmarks to assess the errors. Landmark (.tag) files with clear head registration errors were removed. We observed most errant outlier landmark configurations in the E15.5 and E18.5 embryos, which underwent whole-body registrations. Since the orientation of the head relative to the body cannot be standardized in embryos, the whole-body registrations and inherent constraints of spatial normalization resulted in local registrations errors if their orientation was markedly different from the atlas.

Eliminating problematic outliers with distance distributions is a global solution but not always a local one. For example, if a landmark configuration hardly deviates from the mean on average, yet still has several landmarks with high detection errors, its distance to the mean could be small but its shape distinct. We performed a Principal Component Analysis on each stage-specific landmark dataset (Fig. S2 and S3) to identify such localized errors, assuming the first PC would capture distinctly problematic shapes. Fig. 6 shows the resulting shape distributions along PC1 for each stage. Here, we morphed a surface of the mean shape to each extreme via thin-plate spline and visualized the outputs. If the deformed surface was unusual, we displayed the image and landmarks as above, removed the errant landmark (.tag) file if necessary, and repeated this process until the prediction was correct.

**Figure 6.**
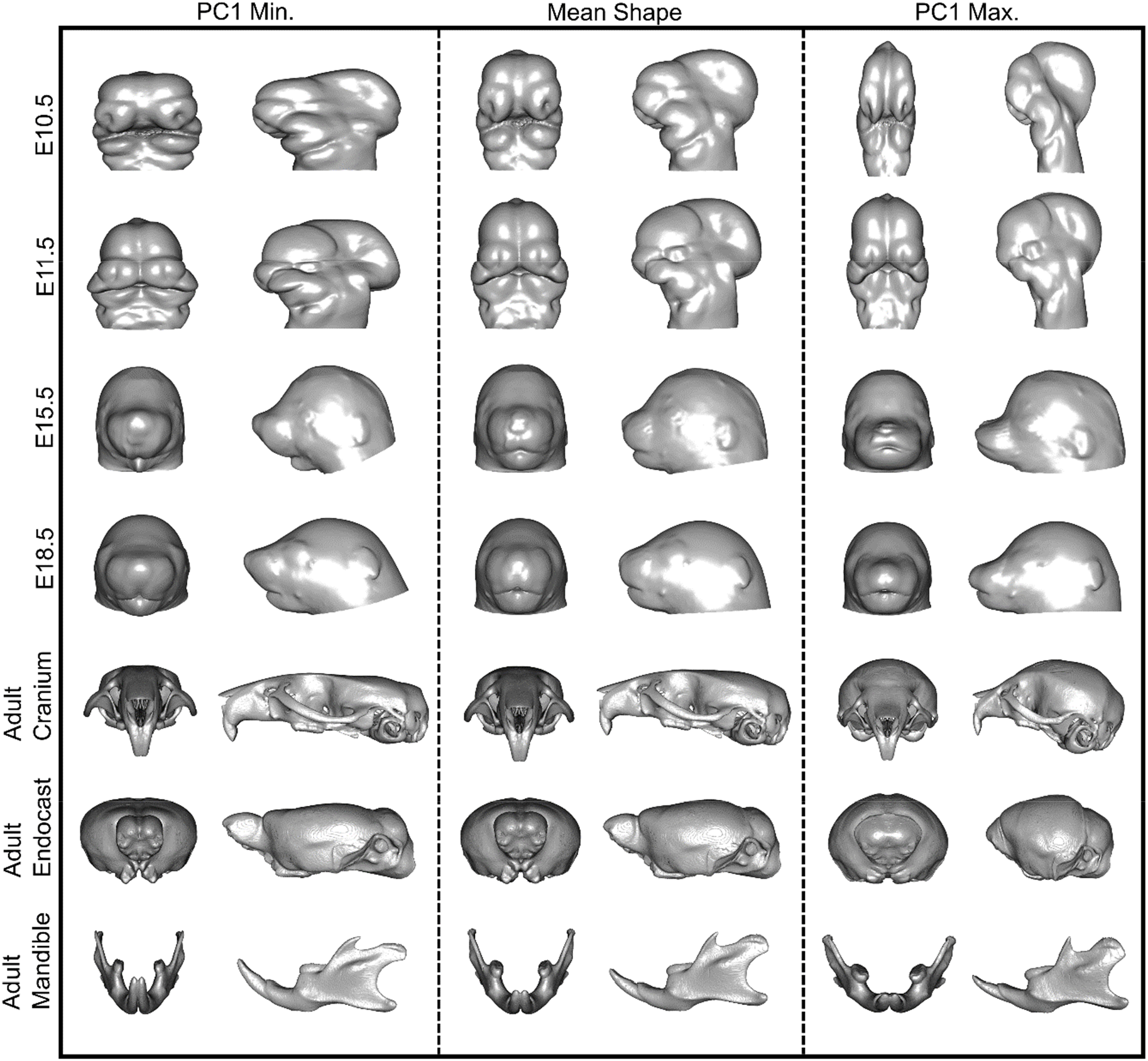
Principal Component Analysis of stage-specific shape data. The mean shape (center) was deformed to the minimum (left) and maximum (right) extremes of PC1. Every morph is shown with anterior and lateral views. Each row represents a different developmental stage, ranging from E10.5 to adulthood.

## Discussion

### Why MusMorph?

The goal of MusMorph was to create a database of standardized mouse morphology data using an automated, high-throughput, and open-source phenotyping pipeline. By combining developmental atlases with a registration and deep learning framework, we constructed common coordinate systems into which various phenotypic data can be integrated. We primarily focused on acquiring morphological data, including anatomical landmarks, segmentations, and deformation fields, for the craniofacial complex and brain. However, we also generated whole body data for other integrative analyses of late-gestation embryos. To enable novel morphometric analyses of genotype-phenotype maps, we utilized mouse models with substantial developmental and genetic variation. Paired alongside other key metadata, such as strain and sex, MusMorph provides the community with a unique opportunity to disentangle the mechanistic basis for morphological variation.

While sparse landmarks are invaluable for geometric morphometrics, there are scenarios where local shape change can be poorly represented. More ambiguous anatomy, such as curves and surfaces, cannot be sufficiently captured with fixed anatomical landmarks, and semi-landmarking each specimen can be tedious and error-prone. Our standardized sparse and dense landmark datasets can enable global and local shape analyses^102,103^, an area in geometric morphometrics historically overlooked. Homologous dense landmark patches across the embryo datasets will also permit joint superimposition of multiple stages into a common shape space for increased statistical power as well as analyses of ontogeny (Fig. S4). In addition to landmarks, we make the corresponding deformation fields available on an ad hoc basis to support voxel-based meta-analyses of morphology. Despite its ubiquitous application in neuroimaging, voxel-based morphometry is rarely seen in fields that study hard tissue, such as evolutionary developmental biology, anthropology, and paleontology. These deformation fields will let one examine internal and external tissue interactions within anatomical context. Finally, we include anatomical segmentations for several stages, which can be used to restrict the dimensionality of a voxel-wise analysis, calculate the size (e.g., volume or surface area) of a structure, or perform a surface-based morphometry analysis. If users are dissatisfied with the coverage of existing landmarks and segmentations, they can modify the atlases and use the image transformations to generate new labels.

We have made the data and scripts freely available at FaceBase (www.facebase.org, doi.org/10.25550/3-HXMC)^35^ and GitHub (https://github.com/jaydevine/MusMorph) to promote transparency, reproducibility, and future data aggregation. Completely open-source efforts like MusMorph are critical for standardizing phenotypic datasets. Unlike the field of genomics, which has been revolutionized through standardized sequencing and data crowdsourcing, phenomics continues to be limited by one-off, self-contained studies that cannot be related to one another. Standardized morphological datasets will allow research groups to, for instance, investigate the effects of a gene mutation alongside other mutants or wildtype strains in a common morphospace. The same can be said for other significant morphological factors and covariates, such as sex and age. Common morphospaces will further encourage multimodal data integration across the phenomic hierarchy, ranging from cellular and developmental phenotyping with light sheet microscopy^104^ to tissue phenotyping with magnetic resonance imaging and contrast-enhanced computed tomography^38^. Large phenotypic datasets will ultimately give us the statistical power needed to interrogate mechanisms that bias and generate morphological variation.

### Sources of error and potential limitations

Staining artifacts are a drawback of contrast-enhanced computed tomography. Among the largest sources of registration error were poor contrast and background noise, particularly in the E15.5 dataset. Variable strain penetrance and inadequate contrast can underrepresent anatomy, whereas background noise can masquerade as anatomy and deceive the registration, even if the alignment is constrained with a mask. We mitigated labelling errors by registering thresholded images and by employing other preprocessing techniques, such as intensity bias correction and normalization. However, in some cases, the intensities of the scanning tube could not be distinguished from the specimen, leading to surface landmark errors. Another spatial alignment problem that was difficult to reconcile was variation in articulated anatomical positions. For example, head orientation relative to the body varied widely among the E15.5 and E18.5 datasets, and mandible orientation relative to the skull differed across the adult dataset. We chose to register the entire scan instead of separate segmentations, masks or cropped volumes, because a) a single registration field is computationally more feasible to generate, store, and use downstream and b) a single atlas with a detailed set of labels is better for data standardization.

Non-linear alignment and labelling errors may occur around extreme anatomical points with high variability. To demonstrate how automated landmark error can be reduced, we implemented a neural network that minimized automated and manual craniofacial shape differences. Since the endocast, mandible, and embryo datasets do not have manual landmark training data, they cannot be optimized. However, if other investigators have training data, a network could be built to correct sparse phenotyping errors in areas of high morphological variability. Lastly, it is important to consider the computational time and memory needed for volumetric registration. To integrate new data, we strongly encourage users to parallelize their work on compute clusters.

### Future development

The majority of MusMorph is composed of head data, because we had reservations about registering whole body data. Now that we have observed no significant differences in registration quality among the datasets, we plan to experiment with more whole-body data for embryos and adults. Another area we intend to improve is our developmental coverage. Despite sampling across most of development, we recognize that additional embryo timepoints (e.g., E9.5 and E12.5-14.5) are needed, as are higher sample sizes throughout mid-gestation and early adulthood. The developing mouse craniofacial complex, for example, undergoes immense growth during the first 30 days after birth^105^. Early postnatal datasets will be critical for asking questions about size and ontogenetic allometry. Finally, to complement our large sample of homozygous embryo mutants, we hope to introduce more wildtype and heterozygous embryos for analyses of normal variation. Heterozygotes have not been a focus of the IMPC, so there is ample opportunity to reveal previously unrecognized embryo phenotypes with standardized MusMorph comparisons. The adult dataset, by contrast, needs to be balanced with more homozygous mutants to better understand how mutations of large effect influence morphological variance and other related phenomena, such as integration and modularity.

## Usage Notes

MusMorph is categorized as a “Project” on FaceBase. Projects can be found in the “Data Browser: Projects” tab at the top of the home page. Project data are organized hierarchically. The levels of the hierarchy in ascending order of data specificity are “Project”, “Dataset”, “Experiment”, and “Biosample”. A project contains datasets, which are sets of similar studies. Each dataset is annotated with study abstracts, experimental designs, and metadata identifiers. Datasets are composed of experiments. An experiment represents a set of similar specimens, so mice with the same genetic background, age, treatment, and mutation would constitute one experiment. Experiments contain biosamples. A biosample is an individual specimen.

After creating a free account and logging in the MusMorph data and metadata can be downloaded at any level in the project hierarchy using the “Export: BDBag” tool at the top-right of the browser. This export function uses DERIVA^106^, the software platform that powers FaceBase, to generate a BDBag (Big Data Bag)^107^ ZIP file. Users then need to download the file and process it via BDBag client tools, either via the command line or GUI application. Specific details about the DERIVA Client installation and the step-by-step export instructions are available here: www.facebase.org/help/exporting.

## Code Availability

Our code is freely available at https://github.com/jaydevine/MusMorph. The scripts describe every stage of the MusMorph data acquisition and analysis, including image preprocessing (e.g., file conversion, image resampling and intensity correction), processing (e.g., atlas generation, non-linear registration, label propagation), and postprocessing (e.g., shape optimization, morphometric analysis). We developed and implemented the code with Bash 4.4.20, R 3.6.1, Python 3.6, and Julia 1.2.0 on Ubuntu. All code is distributed under the GNU General Public License v3.0.

## Acknowledgements

We thank the Advanced Research Computing team at the University of Calgary for facilitating image processing and storage on the ARC and Helix compute clusters. We also thank FaceBase and the International Mouse Phenotyping Consortium for assisting with image data storage and acquisition. Finally, we would like to acknowledge funding from a CIHR Foundation Grant (#159920), an NSERC Discovery Grant (#238992-17), an NIH R01 (#2R01DE019638), an NIH U01 (#U01DE028729), Alberta Innovates, and the Alberta Children’s Hospital Research Institute.

## Author contributions

J.D., M.V.G., and B.H.: Study design, image processing, data collection, data analysis, drafting the manuscript, and revising it critically. H.W. and R.S.M.: Study design, data collection, drafting the manuscript, and revising it critically. W.L., A.N., and L.D.L.: Image processing, data collection, and revising the manuscript critically. R.E.S and A.B.: Data upload and organization. R.M.G., H.A.R., M.M., C.M.U., A.C.N., N.M.Y., P.N.G., C.R., C.J.P., T.W., L.N., A.L.C., A.D.L, A.V., F.R.J., J.M.C., O.K., R.Y.B., A.E.M., R.R.A., D.G., W.D., B.R., M.H., and S.A.M.: Data collection and revising the manuscript critically. All authors gave final approval for publication.

