## Supplementary File for "MusMorph, a database of standardized mouse morphology data for morphometric meta-analyses"

**Table S1.** Summary of MusMorph datasets on FaceBase. Dataset is the name of the dataset. Stage(s) are the development stages within the dataset. N is the sample size. Strain(s) and Genotype(s) are the number of unique strains and genotypes. +/+, +/-, -/-, Other, and Unknown are the number of wildtype, heterozygous, homozygous, other (e.g., neo/null), and unknown specimen zygosities, respectively. F, M, and Unknown are the number of females, males, and unknown sexes. FaceBase DOI is the individual dataset DOI.

| | | | | | Zygosity | | | | | Sex | | | |
|---|---|---|---|---|---|---|---|---|---|---|---|---|---|
| **Dataset** | **Stage(s)** | **N** | **Strain(s)** | **Genotype(s)** | **+/+** | **+/-** | **-/-** | **Other** | **Unknown** | **F** | **M** | **Unknown** | **FaceBase DOI** |
| Ap2 | E10.5, E11.5 | 125 | 1 | 5 | 54 | × | × | 68 | 3 | × | × | 125 | doi.org/10.25550/3-JQMG |
| B9d | E11.5, Adult | 132 | 1 | 4 | 17 | 63 | 31 | × | 21 | 19 | × | 113 | doi.org/10.25550/3-JQMM |
| Nosip | E10.5, Adult | 40 | 1 | 4 | 20 | 13 | 6 | × | 1 | 19 | 10 | 11 | doi.org/10.25550/3-JQMP |
| Mks | E10.5, E11.5 | 15 | 1 | 3 | 6 | 6 | 3 | × | × | × | × | 15 | doi.org/10.25550/3-JVDW |
| Ift | E10.5, Adult | 147 | 2 | 4 | 38 | 103 | 1 | × | 5 | 52 | × | 95 | doi.org/10.25550/3-JVE0 |
| Tctn | E10.5, E11.5 | 51 | 2 | 8 | 8 | 24 | 11 | × | 8 | × | × | 51 | doi.org/10.25550/3-JVE2 |
| Shh | E11.5, Adult | 219 | 1 | 5 | 92 | 125 | × | × | 2 | 36 | 65 | 118 | doi.org/10.25550/3-JVE6 |
| Fgf | E10.5, E11.5, Adult | 250 | 1 | 8 | 95 | 90 | 23 | × | 42 | 24 | 33 | 193 | doi.org/10.25550/3-JVEM |
| Bulgy | E10.5, E11.5, Adult | 262 | 1 | 4 | 99 | 139 | 1 | × | 23 | 105 | 71 | 86 | doi.org/10.25550/3-JZ9G |
| Strain Comparison | E11.5, Adult | 244 | 10 | 9 | 219 | × | × | × | 25 | 103 | 107 | 34 | doi.org/10.25550/3-JZ9J |
| Wnt | E10.5, E11.5 | 136 | 4 | 1 | × | × | × | × | 136 | × | × | 136 | doi.org/10.25550/3-JZ9Y |
| Placenta | E14.5 | 84 | 1 | 8 | 33 | 20 | 31 | × | × | 38 | 24 | 22 | Unreleased |
| IMPC | E15.5, E18.5 | 3001 | 1 | 419 | 881 | 36 | 2084 | × | × | 1387 | 1418 | 196 | doi.org/10.25550/3-JZA6 |

| | | | | | | | | | | | | | |
|---|---|---|---|---|---|---|---|---|---|---|---|---|---|
| Spry | Adult | 260 | 3 | 9 | 88 | 114 | 58 | × | × | 108 | 152 | × | doi.org/10.25550/3-JZAM |
| Osteo Imperfecta | Adult | 29 | 1 | 2 | 29 | × | × | × | × | 16 | 13 | × | doi.org/10.25550/3-KB00 |
| Enhancer | Adult | 465 | 3 | 17 | 138 | 55 | 270 | × | × | 208 | 254 | 3 | doi.org/10.25550/3-KB02 |
| Ghrhr | Adult | 394 | 1 | 2 | × | 105 | 289 | × | × | 57 | 87 | 250 | doi.org/10.25550/3-KB08 |
| Nipbl | Adult | 59 | 1 | 2 | 37 | 22 | × | × | × | 10 | 49 | × | doi.org/10.25550/3-KB0J |
| Collaborative Cross | Adult | 1129 | 60 | 58 | 1129 | × | × | × | × | 538 | 591 | × | doi.org/10.25550/3-KB0W |
| BBDS | Adult | 24 | 1 | 5 | 6 | 11 | 7 | × | × | 17 | 7 | × | Unreleased |
| Brachymorph | Adult | 30 | 1 | 1 | × | × | 30 | × | × | 14 | 10 | 6 | doi.org/10.25550/3-KB1W |
| Hybrid | Adult | 817 | 20 | 19 | 817 | × | × | × | × | 444 | 373 | × | doi.org/10.25550/3-KB32 |
| Brain-Face | Adult | 141 | 2 | 6 | × | 40 | 74 | 26 | 1 | 75 | 66 | | doi.org/10.25550/3-KB3J |
| RASopathy | Adult | 39 | 1 | 2 | 21 | | 18 | | | 20 | 19 | | Unreleased |
| Bmp | Adult | 274 | 1 | 24 | 115 | 131 | 26 | 2 | × | 121 | 150 | 3 | doi.org/10.25550/3-KB46 |
| Lrp | Adult | 9 | 1 | 1 | × | 9 | × | × | × | 9 | × | × | doi.org/10.25550/3-KB4J |
| Diversity Outbred | Adult | 1048 | 1 | 8 | 1048 | × | × | × | × | 574 | 337 | 137 | doi.org/10.25550/3-KB4P |
| Longshanks | Adult | 446 | 1 | 5 | 446 | × | × | × | × | 246 | 200 | × | doi.org/10.25550/3-KFBE |
| MPS | Adult | 45 | 1 | 1 | × | × | 45 | × | × | 12 | 28 | 5 | doi.org/10.25550/3-KFBY |
| Nabo | Adult | 90 | 1 | 2 | × | 43 | 47 | × | × | 39 | 51 | × | Unreleased |
| Pten | Adult | 26 | 1 | 2 | 2 | × | 24 | × | × | 12 | 14 | × | doi.org/10.25550/3-KFZJ |
| Trp | Adult | 25 | 1 | 1 | × | 25 | × | × | × | 23 | 2 | × | doi.org/10.25550/3-KFZW |

**Table S2.** Sparse embryo anatomical landmarks/derivatives and their definitions (* indicates landmarks that are specific to the E14.5, E15.5, and E18.5 embryos).

| **Paired Landmarks (R/L)** | **Anatomical Definition** |
|---|---|
| 1/2 | Caudal most junction of the lateral nasal process and maxillary process |
| 3/4 | Nasal aperture and rostral ventral most junction of the lateral nasal process and maxillary process |
| 5/6 | Corner of the mouth |
| 7/8 | Medial, rostral, dorsal corner of the mandibular process |
| 11/13 | Dorso-caudal most point of the lateral nasal process |
| 12/14 | Center of the eye |
| 15/16 | Dorsal most point of the nasal aperture |
| 21/22 | Junction between the bulge of the trigeminal ganglion and pontine flexure of the developing brain |
| 23/24* | Dorso-caudal corner of whisker row |
| **Midline Landmarks** | **Anatomical Definition** |
| 9 | Rostral midline point of the mandibular processes |
| 10 | Rostral most point at the midline of the medial nasal processes. |
| 17 | Midline dorsal most extent of the face |
| 18 | Dorsal midline junction between the growing forebrain and midbrain lobes |
| 19 | Dorso-caudal most point on the midline of the midbrain |
| 20 | Caudal most midline point at the back of the head, just ventral to the midbrain |
| 25* | Tip of the nose and rostral most point of the face |

**Table S3.** Embryo landmark patch vertices, their size, the number of semilandmarks, and their row position in an individual array. Due to the extra sparse landmarks for E14.5-15.5 and E18.5, their patch positions are given in parentheses "()".

| Vertex 1 | Vertex 2 | Vertex 3 | Patch Size | No. of Semis | Rows in Array |
|---|---|---|---|---|---|
| 20 | 19 | 21 | 10 | 36 | 23:58 (26:61) |
| 20 | 19 | 22 | 10 | 36 | 59:94 (62:97) |
| 18 | 19 | 21 | 10 | 36 | 95:130 (98:133) |
| 18 | 19 | 22 | 10 | 36 | 131:166 (134:169) |
| 18 | 14 | 22 | 10 | 36 | 167:202 (170:205) |
| 18 | 12 | 21 | 10 | 36 | 203:238 (206:241) |
| 5 | 12 | 21 | 10 | 36 | 239:274 (242:277) |
| 17 | 12 | 18 | 10 | 36 | 275:310 (278:313) |
| 17 | 14 | 18 | 10 | 36 | 311:346 (314:349) |
| 6 | 14 | 22 | 10 | 36 | 347:382 (350:385) |
| 6 | 14 | 16 | 7 | 15 | 383:397 (386:400) |
| 5 | 12 | 15 | 7 | 15 | 398:412 (401:415) |
| 5 | 7 | 9 | 5 | 6 | 413:418 (416:421) |
| 6 | 8 | 9 | 5 | 6 | 419:424 (422:427) |
| 13 | 16 | 17 | 5 | 6 | 425:430 (428:433) |
| 11 | 15 | 17 | 5 | 6 | 431:436 (434:439) |
| 15 | 16 | 17 | 4 | 3 | 437:439 (440:442) |
| 3 | 10 | 15 | 4 | 3 | 440:442 (443:445) |
| 4 | 10 | 16 | 4 | 3 | 443:445 (446:448) |
| 7 | 8 | 9 | 4 | 3 | 446:448 (449:451) |
| 10 | 15 | 16 | 4 | 3 | 449:451 (452:454) |

**Table S4.** Sparse adult craniofacial landmarks and their definitions.

| Paired Landmarks (R/L) | Anatomical Definition |
|---|---|
| 2/1 | Superior point of post-tympanic hook |
| 4/3 | Paroccipital process |
| 6/5 | Posterior point on internal pterygoid process |
| 14/13 | Lateral point on frontal suture |
| 16/15 | Lateral zygomatic-frontal suture |
| 18/17 | Posterior zygomaticofrontal junction |
| 20/19 | Posterior margin of malar process |
| 22/21 | Frontal-temporal-parietal junction |
| 23/24 | Anterior margin of incisive foramen |
| 25/26 | Medial maxilla-premaxilla junction |
| 27/28 | Anterior inferior zygomatic |
| 29/30 | Anterior temporo-zygomatic junction |
| 31/32 | Anterior superior alveoli |
| 33/34 | Posterior incisive foramen |
| 35/36 | Point along palatine-maxillary suture |
| 37/38 | Medial palatal-pterygoid junction |
| 39/40 | Posterior superior alveoli |
| 41/42 | Lateral palatal-pterygoid junction |
| 43/44 | Spheno-occipital synchondrosis |
| 45/46 | Anterior foramen ovale |
| 47/48 | Posterior temporo-zygomatic junction |
| 49/50 | Auditory-temporal-sphenoid junction |
| 51/52 | Anterior inferior auditory bulla |
| 53/54 | Occipital-auditory-sphenoid junction |
| 55/56 | Point along occipitomastoid suture |
| 57/58 | Medial occipital condyle |
| 59/60 | Anterior nasal and premaxilla |
| 61/68 | Frontal suture on orbital rim |
| 62/69 | Superior temporo-zygomatic suture |
| 63/70 | Posterior zygomatic process |
| 64/71 | Superior posterior tympanic ring |
| 65/72 | Occipital-auditory junction |
| 66/67 | Midline superior incisor |
| 91/79 | Posterior tympanic ring |
| 80/81 | Anterior inferior maxilla |
| 82/85 | Medial point of first upper molar |
| 83/86 | Medial point of second upper molar |
| 84/87 | Medial point of third upper molar |

| | |
|---|---|
| 89/90 | Superior lateral point of paroccipital process |
| 92/93 | Superior lateral most point of occipital condyle |
| **Midline Landmarks** | **Anatomical Definition** |
| 7 | Posterior point of presphenoid |
| 8 | Superior most point of foramen magnum |
| 9 | Posterior most point of occipital |
| 10 | Lambda |
| 11 | Bregma |
| 12 | Nasion |
| 73 | Anterior foramen magnum |
| 74 | Midline junction between the basioccipital and sphenoid |
| 75 | Midline junction between the sphenoid and presphenoid |
| 76 | Anterior junction of the endocranial presphenoid |
| 77 | Endocranial junction between the frontal and ethmoid |
| 78 | Anterior most point of nasal bone |
| 88 | Anterior point on alveolar process between incisors |

**Table S5.** Sparse adult endocast landmarks and their definitions.

| Paired Landmarks (R/L) | Anatomical Definition |
|---|---|
| 5/4 | Trigeminal nerve |
| 7/6 | Lateral junction between cerebellum and medulla |
| 9/8 | Lateral junction between occipital lobe and cerebellum |
| **Midline Landmarks** | **Anatomical Definition** |
| 1 | Distal most point of olfactory bulb |
| 2 | Midline junction between olfactory bulb and anterior olfactory nucleus |
| 3 | Midline junction between anterior olfactory nucleus and ventral striatum, optic nerves |
| 10 | Midline junction between midbrain and cerebral cortex |
| 11 | Midline junction between olfactory bulb and cerebral cortex |
| 12 | Optic chiasma |

**Table S6.** Sparse adult mandible landmarks and their definitions.

| **Paired Landmarks (R/L)** | **Anatomical Definition** |
|---|---|
| 1/6 | Posterior mandibular angle |
| 2/7 | Posterior mandibular condyle |
| 3/8 | Posterior superior most point of coronoid process |
| 4/9 | Mandibular tuberosity and posterior point of molar alveolar rim |
| 5/10 | Anterior point of molar alveolar rim |
| 13/14 | Inferior most point of mental protuberance |
| 15/18 | Mandibular foramen/inferior alveolar foramen |
| 16/17 | Superior anterior most point of molar row |
| **Midline Landmarks** | **Anatomical Definition** |
| 11 | Mental spine and posterior midline bone-tooth junction |
| 12 | Anterior superior most point of incisor alveolar rim |
| 19 | Anterior most midline point of incisors |

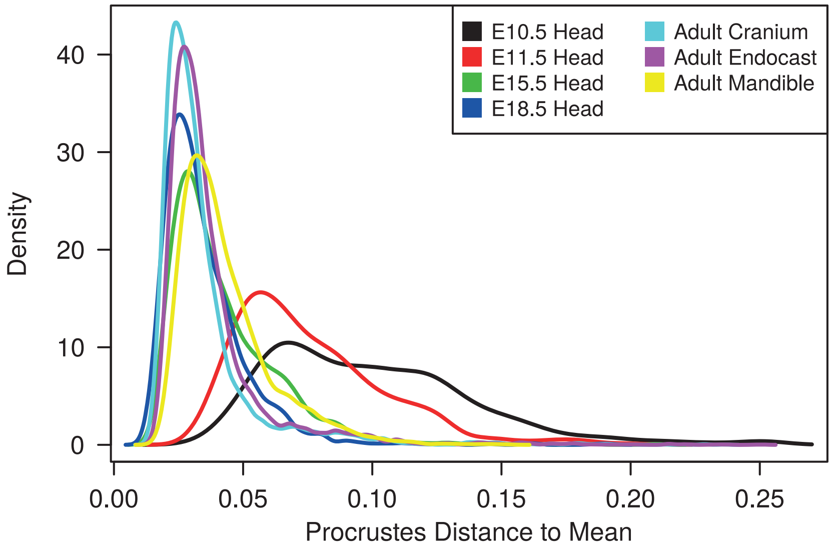

**Figure S1.** Density plot of Procrustes distances to the mean shape for each stage. These distributions were used to identify and eliminate global outliers.

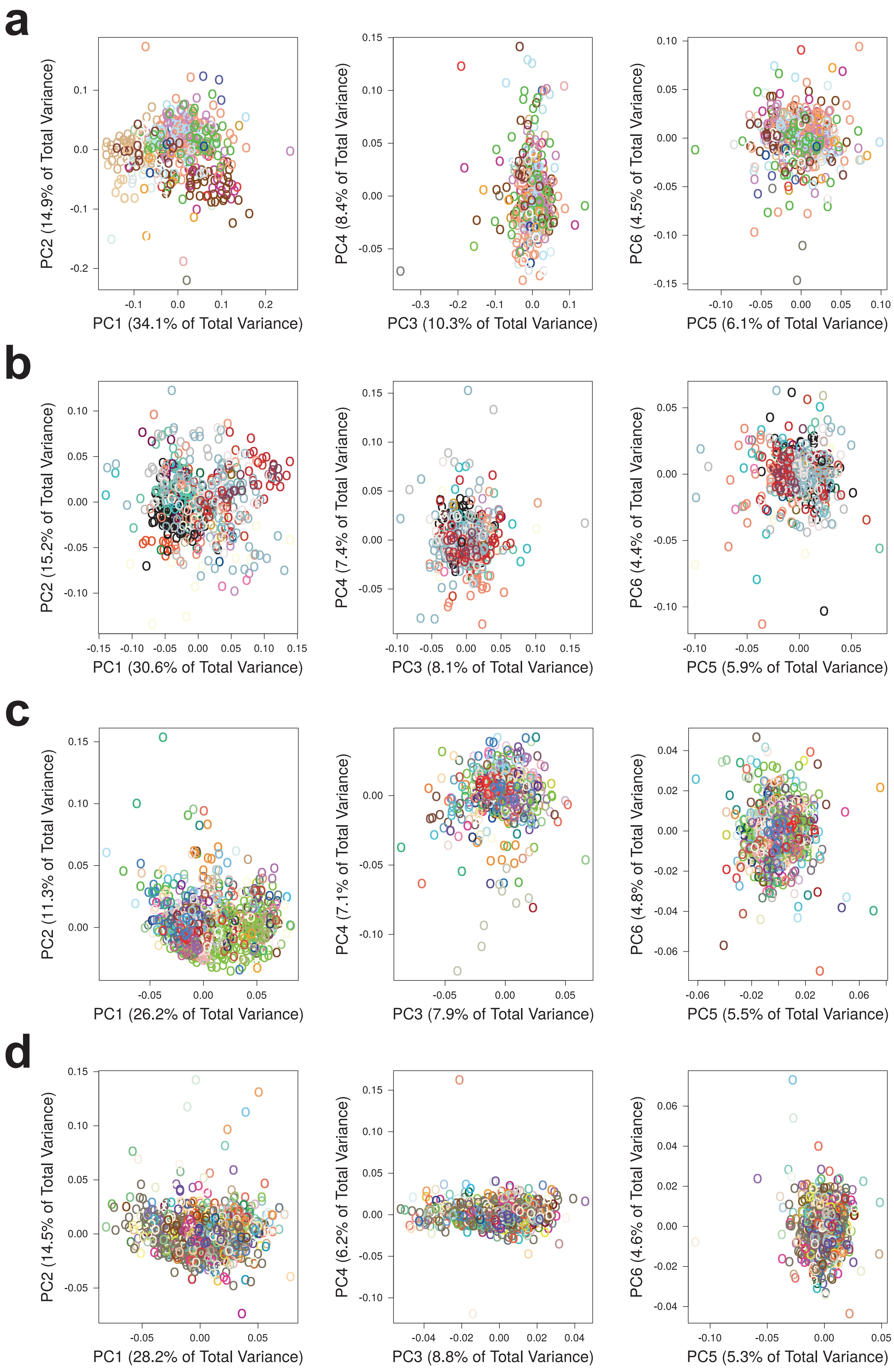

**Figure S2.** Principal Component Analysis of embryo landmark data. The first six PCs for (a) E10.5, (b) E11.5, (c) E15.5, and (d) E18.5 are shown. Each color represents a unique genotype within the stage. These data were used to identify and eliminate local outliers, as well as visualize stage-specific shape distributions.

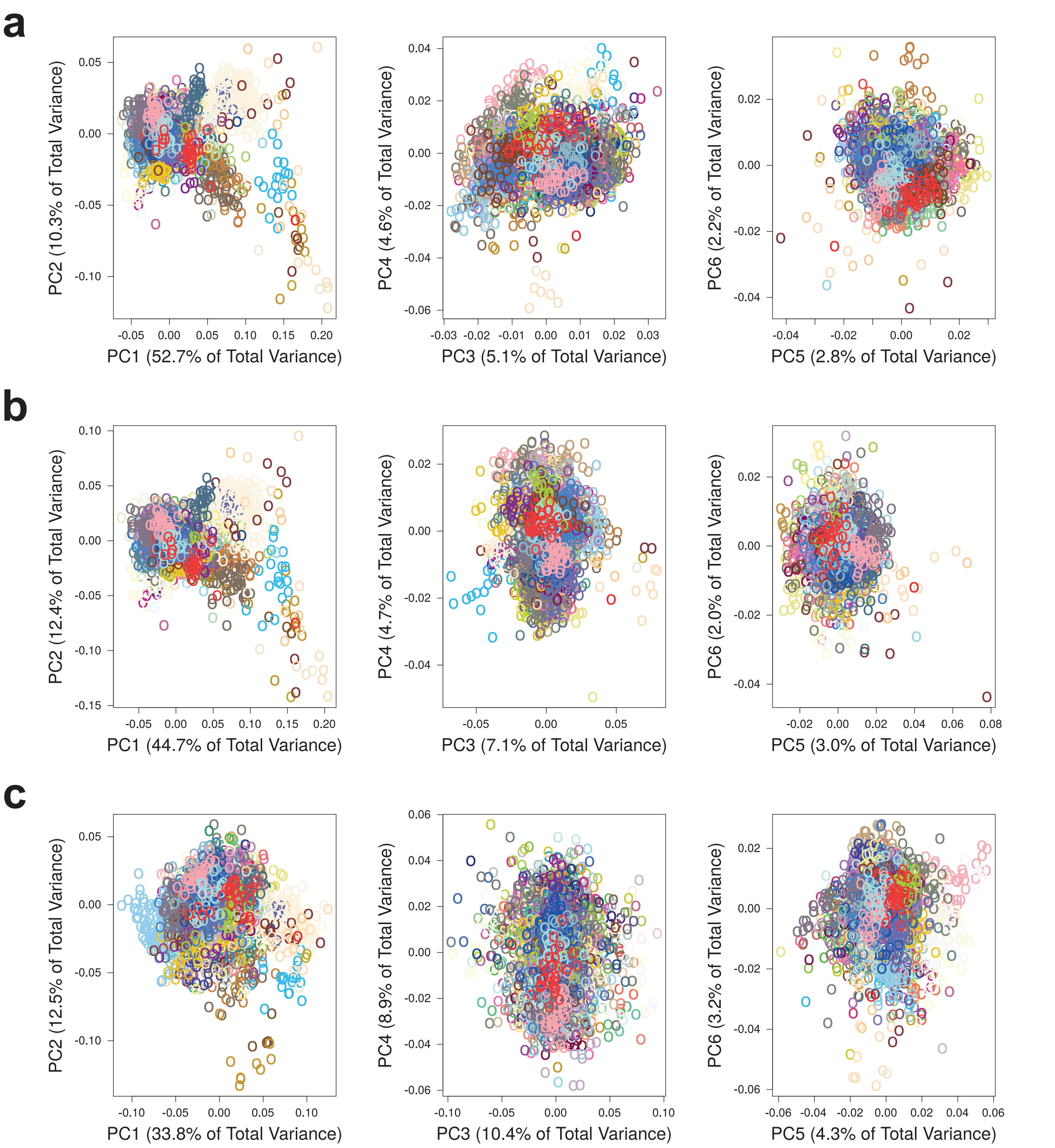

**Figure S3.** Principal Component Analysis of adult landmark data. The first six PCs for the (a) cranium, (b) endocast, and (c) mandible are shown. Each color represents a unique genotype. These data were used to identify and eliminate local outliers, as well as visualize the stage-specific shape distributions.

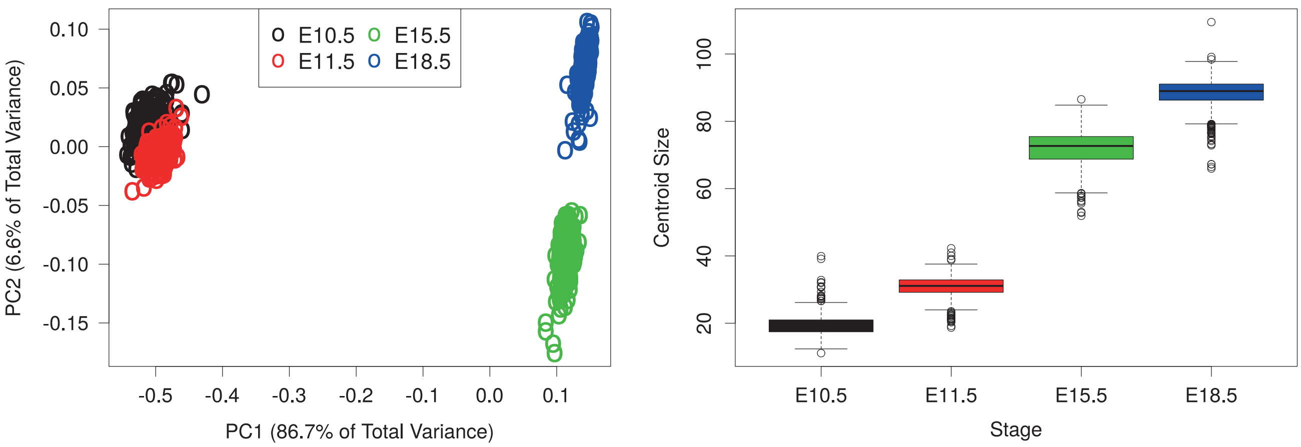

**Figure S4**. Generalized Procrustes Analysis of all embryo stages. For the sake of demonstration, an equivalent number of specimens were sampled from each stage (N=500) to not bias the mean. Left: Principal Component Analysis showing the large allometric effect along PC1. Right: Boxplots of centroid size obtained from the Procrustes superimposition.
